# Varenicline rescues nicotine-induced decrease in motivation for sucrose reinforcement

**DOI:** 10.1101/2020.06.03.129791

**Authors:** Erin Hart, Daniel Hertia, Scott T Barrett, Sergios Charntikov

## Abstract

Varenicline is one of the top medications used for smoking cessation and is often prescribed before termination of nicotine use. The effect of this combined nicotine and varenicline use on the reward system and motivation for primary reinforcement is underexplored. The goal of this study was to assess the effects of nicotine and varenicline on the consumption of sucrose. In Experiment 1, we first assessed the responding for sucrose after pretreatment with nicotine (0, 0.1, or 0.4 mg/kg) and varenicline (0.0, 0.1, 1.0 mg/kg) using a behavioral economics approach. The responding for sucrose was then assessed using a progressive ratio schedule of reinforcement after pretreatment with all possible combinations of nicotine and varenicline doses. In Experiment 2, rats were assessed for the consumption of sucrose in home cages after pretreatment with nicotine and varenicline. We found that a) nicotine decreased economic demand for sucrose, b) varenicline rescued nicotine-induced reduction in economic demand for sucrose, and c) history of varenicline treatment predicted responding for sucrose on a progressive ratio schedule of reinforcement where rats with a history of varenicline treatment responded significantly lower for sucrose across nicotine doses than rats that have not being exposed to varenicline. The results of Experiment 2 largely confirmed that nicotine decreases motivation for sucrose using a passive consumption protocol and that varenicline rescues this effect. Overall, these findings suggest that varenicline interacts with the effects of nicotine by restoring nicotine-induced reduction in motivation for appetitive rewards.

## 1. Introduction

Global use of nicotine products is responsible for over 7 million deaths each year and is the leading cause of preventable deaths worldwide (World Health Organization, 2017). The annual economic burden of nicotine use in the US alone is around $400 billion dollars (USDHHS, 2014). There are many users that desire to quit their nicotine habit, but few are successful in doing so (70-80 % relapse rate with intervention; Cahill et al., 2013). Varenicline (trade name Chantix and Champix) is one of the top three pharmacological treatments prescribed for the cessation of nicotine use. Treatment with varenicline is marginally more effective than placebo and increases cessation rates by approximately 10-17 % over placebo depending on the study and outcome criteria (Cahill et al., 2013; Rosen et al., 2018). Varenicline is marketed as a treatment offering flexibility in setting a quit date (Pfizer, 2020). Patients prescribed varenicline usually take varenicline while continuing smoking for 1 to 12 weeks with additional varenicline treatment thereafter (up to additional 12 weeks; Tonstad et al., 2006; Pfizer, 2020). The effect of this regiment on primary reinforcers is underexplored. Therefore, this study was designed to start filling a gap in our understanding regarding how varenicline and nicotine interact to affect food reward in a preclinical setting.

Nicotine is a stimulant and a naturally occurring insecticide in the nightshade family of plants. Nicotine is a mild reinforcer when compared to other stimulants like cocaine or methamphetamine (Dougherty et al., 1981; Le Foll and Goldberg, 2009). Indeed, responding for nicotine in preclinical self-administration studies is only marginally higher than responding for saline and is often insensitive to variation in nicotine dose (Huynh et al., 2017; Kohut and Bergman, 2016). The fact that nicotine is a weak primary reinforcer is somewhat incongruent with the fact that it is one of the most commonly abused substances. The wide use of nicotine products has been attributed to the complex nature of nicotine reward that involves genetic, biological, learning, and pharmacological dimensions, to name a few (Donny et al., 2011; Dougherty et al., 1981; Garcia-Rivas and Deroche-Gamonet, 2019). Contingent or non-contingent administration of nicotine can enhance the rewarding effects of other non-pharmacological stimuli (Chaudhri et al., 2006; Donny et al., 2003; Liu et al., 2007; Palmatier et al., 2007). For example, rats will increase their responding for a visual stimulus independently of whether nicotine is self-administered or administered by the experimenter (Donny et al., 2003). Furthermore, nicotine can amplify the incentive salience of other reward-associated stimuli, which may promote increased nicotine use and escalate the development of nicotine dependence (Palmatier et al., 2013). Finally, nicotine can serve as an interoceptive stimulus and can acquire conditioned reinforcing properties when it is paired with an appetitive reward like sucrose (Bevins et al., 2012; Charntikov et al., 2020). Specifically, rats will work harder for nicotine on a progressive ratio schedule of reinforcement after a history of nicotine-sucrose pairings, indicating that nicotine can acquire additional reinforcing properties through associations with other rewards (Charntikov et al., 2020). Thus the high abuse prevalence of tobacco products likely stems from the multifaceted nature of nicotine reward at the junction of environmental, biological, and genetic factors.

Weight gain is a significant concern among those considering smoking cessation (Hall et al., 1986; Perkins, 1993). On average, smokers weigh less than non-smokers and tend to gain weight after cessation of smoking (Albanes et al., 1987; Froom et al., 1998; Hall et al., 1986; Jacobs and Gottenborg, 1981; Klesges et al., 1997; Perkins, 1993). However, human studies do not show a decrease in food intake among smokers. That is, there is no effect of acute or chronic nicotine use on food consumption in individuals maintaining a regular smoking regiment (Perkins, 1992). In rats, on the other hand, nicotine produces anorectic effects resulting in a reduction of food intake (Grebenstein et al., 2013; Grunberg et al., 1988; Miyata et al., 2001; Winders and Grunberg, 1990). For example, contingent or non-contingent nicotine administration reduces normal weight gain in rats (Grebenstein et al., 2013; Rupprecht et al., 2016). The effect of nicotine on food consumption in preclinical studies is inconsistent, with some studies showing no effect on food intake (e.g., Rupprecht et al., 2016) while other studies showing a decrease in food consumption (e.g., Grebenstein et al., 2013; for review see Donny et al., 2011). Food is a primary reinforcer and the reinforcing salience of foods is an important factor in food consumption (Epstein and Leddy, 2006). The reinforcement value of a food stimulus may be characterized as the amount of effort an individual is willing to expend to earn access to that food stimulus. To better understand how varenicline impacts the cessation of nicotine use, it is important to understand how nicotine affects work performance for food reinforcement over a range of response requirements. Furthermore, because varenicline is often prescribed to patients while they use nicotine products, it is important to understand how varenicline interacts with nicotine to impact food consumption under a range of controlled experimental conditions.

Varenicline is an *α*4*β*2 nicotinic acetylcholine receptor (nAChR) partial agonist and a full agonist for *α*3*β*2 and *α*7 nAChRs. Varenicline is extensively used around the world as a first treatment option for smoking and is the most effective smoking cessation pharmacotherapeutic currently available (Cahill et al., 2013; Gómez-Coronado et al., 2018). The mechanism by which varenicline improves smoking cessation rates is not fully understood, but the existing evidence shows that the increase of dopamine in the nucleus accumbens is an important factor in reducing cravings and withdrawal effects associated with nicotine use. Acute and chronic varenicline treatment protocols consistently show reduction in nicotine self-administration in preclinical studies (George et al., 2011; O’Connor et al., 2010; Rollema et al., 2007). However, a recent study from our laboratory shows that duration of access to nicotine is associated with the variation in the economic demand for nicotine and response to treatment. Specifically, a long-access self-administration protocol increases economic demand for nicotine, and rats self-administering nicotine for 12 h a day do not show a decrease in nicotine self-administration following pretreatment with varenicline (Kazan et al., 2020). With that in mind, the effectiveness of varenicline in clinical studies may be in part explained by its ability to substitute for the mildly reinforcing or reward-enhancing effects of nicotine. For example, varenicline dose-dependently enhances responding for a moderately reinforcing visual stimulus and is able to attenuate the reinforcement-enhancing effects of nicotine in preclinical studies (Levin et al., 2012; Schassburger et al., 2015). Furthermore, varenicline increases the rewarding effects of sensory reinforcement across a broad range of workload requirements as assessed using the established reinforcer demand model (Barrett et al., 2018). In that study, both nicotine and varenicline (to a lesser extent) enhanced the reinforcement value of a mildly-reinforcing visual stimulus in male and female rats. Because the reinforcer demand approach assesses responding over a range of workload requirements, this approach may provide a more comprehensive view of behavioral change under different pharmacological treatments.

Reinforcer demand modeling is used to study behavioral responses maintained by a variety of reinforcers in both clinical and preclinical studies. The reinforcer demand approach to studying motivation for reinforcers has been adapted from microeconomics theory, which relates the consumption of goods to the consumption expenditure (Hursh and Roma, 2016). This approach has been extensively used to assess behavioral responses for a variety of reinforcers, including sensory stimulation, food, and drugs, to name a few (Hursh et al., 2005; Hursh and Roma, 2016; R. Hursh, 2014). Using this approach, rats can be trained to respond for a reinforcer on a low fixed ratio (FR) schedule of reinforcement which is then gradually increased over successive sessions, resulting in an increase in the “cost” to obtain a reinforcer. Thus, the reinforcer in this setting is conceptualized as a “good,” response output maintained by the reinforcer is conceptualized as “consumption expenditure,” and FR schedule is conceptualized as “cost.” Using this approach, experimenters can assess grouped and individual economic demand for the reinforcer (Kazan and Charntikov, 2019; Kazan et al., 2020; Stafford et al., 2019). Although many parameters can be derived from the behavioral economics model, the strength of the reinforcer represented by the degree to which an animal is willing to work to receive a reinforcer can be distilled into a single variable—*essential value*. The main advantage of using *essential value* is that it is a unifying measure that takes into consideration consumption when the price for a reinforcer is low (e.g., FR1), when the price for the reinforcer is high (e.g., higher or terminal FR schedules), and the slope of the demand curve (elasticity). That is, the *essential value* represents the sensitivity of changes in consumption of a reinforcer to changes in response cost to obtain that reinforcer, across the full range of the demand function. Our current study was designed to assess the effects of varenicline and nicotine on responding for sucrose using behavioral economics approach outlined above.

## 2. Materials and methods

### 2.1. Animals

Subjects were experimentally naïve, male Sprague-Dawley rats (250-300g) purchased from Envigo (Indianapolis, IN, USA). Rats were housed individually in a temperature- and humidity-controlled colony. Experiments were conducted during the light portion of a 12-hour light/dark cycle (lights on at 7 a.m.). Following seven days of acclimation to the colony, rats were handled 2 min/day for 5 consecutive days. Water was freely available; access to chow was restricted after the acclimation period to maintain rats at 85 % of their free-feeding body weight. This 85 % target weight was increased by 2 g every four weeks from the beginning of the study. All procedures were in accordance with the Guide for the Care and Use of Laboratory Animals (National Research Council et al., 2010) and were reviewed and approved by the University of New Hampshire Institutional Animal Care and Use Committee.

### 2.2. Apparatus

Conditioning chambers (ENV-018MD; Med Associates, Inc.; St. Albans, VT, USA; 30.5 × 24.1 × 21.0 cm), were enclosed in a sound- and light-attenuating cubicle equipped with an exhaust fan. Each chamber had aluminum side-walls, metal rod floors with polycarbonate front, back, and ceiling. A recessed receptacle (5.2 × 5.2 × 3.8 cm; l × w × d) was centered on one sidewall. A dipper arm, when raised, provided access to 100 *μ*L of 5 % (w/v) sucrose solution in the receptacle. Med Associates interface and software (Med-PC for Windows, version IV) were used to record data and all programmed events.

### 2.3. Drugs

Nicotine bitartrate (MP Biomedicals; Solon, OH, USA) and varenicline dihydrochloride (a gift from NIDA Drug Supply Program; Bethesda; MD, USA), were dissolved in 0.9 % sterile saline. Doses and administration protocols were adopted from previous research (Charntikov et al., 2017; George et al., 2011; Kazan et al., 2020; Wouda et al., 2011). Nicotine doses are reported as base, whereas varenicline doses are reported as salt.

### 2.4. Preliminary lever training

Rats in Experiments 1 and 2 had experienced an identical lever training protocol, outlined as follows. Rats were first trained to retrieve liquid sucrose (5 % w/v; 100 *μ*L) from a dipper receptacle until reaching an 80% retrieval criterion (3-5 days). These 50-min dipper training sessions consisted of non-contingent sucrose presentations delivered on a variable time schedule (~ 3 rewards per minute). Rats were then trained to lever press for liquid sucrose (5 % w/v; 100 *μ*L). At the start of each session, the house-light was turned on and a randomly selected lever (right or left) was inserted. A lever press or lapse of 15 s resulted in sucrose delivery (4 s access), lever retraction, and commencement of a timeout (average=60 s; range=30 to 89 s). Following the timeout, a randomly selected lever was inserted with the condition that the same lever could not be presented more than twice in a row. This protocol was repeated for 60 sucrose deliveries. Sessions lasted 65 to 80 min depending on individual performance. Training continued until a lever press was made on at least 80 % of the lever insertions for two consecutive days (i.e., 3 to 6 sessions). Rats were then pseudo-randomly assigned an active lever (right or left) so that there was an equal number of right and left active levers in each condition. Rats were then trained to lever press on their active lever using a fixed ratio 1 (FR1) schedule of reinforcement for six consecutive days. Each of these 30-min training sessions began with the termination of the house light and extension of both levers into the conditioning chamber. Following a response on the active lever both levers were retracted and the dipper arm was raised to allow access to liquid sucrose for 10 seconds. Following sucrose presentation, the dipper arm was lowered and both levers were extended back into the chamber.

### 2.5. Experiment 1: The interaction effect of varenicline and nicotine on economic demand for sucrose

Rats were first treated with 3 daily subcutaneous injections of nicotine (0.4 mg/kg) to alleviate aversive effects associated with initial exposure to nicotine. Rats were then randomly assigned to three experimental conditions to form a 3 × 3 mixed-subjects factorial design with Varenicline Condition (Var; 0.0, 0.1, or 1.0 mg/kg) as a between-subjects variable and Nicotine Dose (Nic; 0.0, 0.1, or 0.4 mg/kg) as a within-subjects variable (see Figure 1) for major experimental progression). Prior to daily sucrose self-administration sessions, rats were injected intraperitoneally with the assigned varenicline dose (30 minutes before the start) and then subcutaneously with nicotine dose 5 minutes before the start of each session. Daily nicotine doses were assigned to each rat using Latin square design. Rats were allowed to earn liquid sucrose (5 % w/v; 100 *μ*L) using fixed ratio (FR) schedules of reinforcement that were escalated every three days after completion of testing with each nicotine dose for each FR test block (1, 3, 5, 8, 12, 18, 26, 38, 58, 86, 130, 195, and 292; 30-min session length). This design allowed all rats to earn sucrose on the same schedule under differing nicotine doses until failing to earn at least one infusion on all three days of the same schedule of reinforcement. Rats were then retrained on the FR1 schedule of reinforcement for at least two consecutive daily sessions and then trained on a progressive ratio (PR; 1, 3, 6, 10, 15, 20, 25, 32, 40, 50, 62, 77, 95, 118, 145, 179, 219, 268, and 328) schedule for additional five days prior to advancing to the next phase of this experiment. In the next phase of this experiment, all rats were tested with all possible combinations of varenicline and nicotine doses (3 × 3 factorial design yielding nine total test combinations) using a PR schedule of reinforcement (30-min sessions).

**Figure 1:**
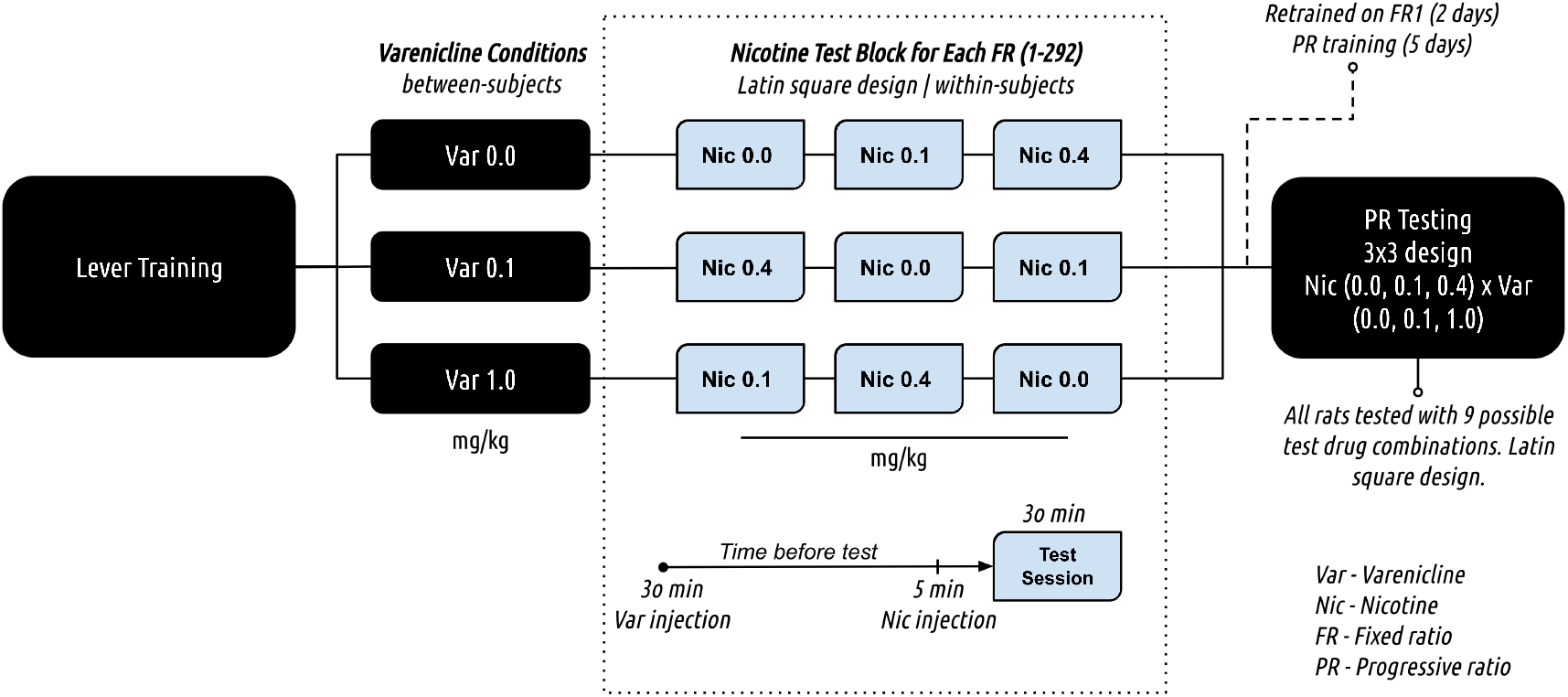
Experimental progression. This experiment used a mixed-subject design with Varenicline Condition as a between-subjects factor and Nicotine Dose as a within-subjects factor. Rats were first trained to press a lever for a reinforcer and then were randomly assigned to one of three Varenicline Conditions. Rats were pretreated with a combination of varenicline (0.0, 0.1, or 1.0 mg/kg) and nicotine (0.0, 0.1, 0.4 mg/kg) prior to each daily session where rats were lever pressing for liquid sucrose on progressively escalated FR schedules of reinforcement. Thereafter, rats were retrained to lever press on FR2 and trained to lever press on PR schedules of reinforcement. During the final phase, all rats were tested with all possible combinations of varenicline and nicotine doses previously used in the experiment.

### 2.6. Experiment 2: The effect of nicotine, varenicline, or their combination on the sucrose consumption in home cages

Rats were lever trained as described above. Rats were then allowed to self-administer sucrose to acquire individual economic demand for sucrose, as described above. Individual consumption of sucrose during the economic demand phase was used to assign subjects to one of 4 experimental conditions ensuring that approximately equal number of rats with higher and lower demands for sucrose were assigned to each group. These 4 experimental conditions were formed using a 2 × 2 between-subjects factorial design with varenicline (Var; 0.0 or 1.0 mg/kg) and nicotine (Nic; 0.0 or 0.4 mg/kg) as independent variables. Doses for this experiment were chosen based on results obtained in Experiment 1 where these doses resulted in statistically detectable effects relevant to this follow-up experiment. After preliminary lever training and the acquisition of economic demand for sucrose, rats were allowed to consume liquid sucrose (5% w/v; 100 mL bottles) in home cages. Prior to this phase, all rats were pretreated with nicotine (0.4 mg/kg; 3 days) as was described above. One hour prior to each daily access to sucrose session, bottles containing water were removed from all home cages. Rats were then injected with the assigned varenicline dose 30-min before—and with the assigned nicotine dose 5 min before—sucrose bottles were made available. At the end of the 30-min consumption period, sucrose bottles were replaced with water bottles and the remaining sucrose solution was measured using a graduated cylinder to estimate individual sucrose consumption.

### 2.7. Dependent measures and statistical analyses

#### 2.7.1. Analytical approach

##### 2.7.1.1. Essential value assessment

The essential value was derived from the economic demand model that was calculated from the nonlinear least squares regression model fit to the individual active lever pressing data from each schedule of reinforcement using the following formula: 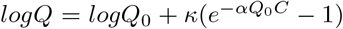. In this formula *Q* represents reinforcer consumption, *Q_0_* is a consumption when the price is zero or free, *κ* is a constant for the range of demand, *e* is the base of the natural logarithm, *C* is the varying cost of each reinforcer, and *α* is the rate of decline in relative *log* consumption with increases in price. The essential value was calculated from the demand model using the following formula *EV* = 1 ÷ (100 × *α* × *κ*^1.5^).

##### 2.7.1.2. Experiment 1

Active lever presses and economic demand for sucrose were used as primary dependent measures. Economic demand parameters were assessed using the demand model by Hursh and Silberberg (Hursh and Silberberg, 2008; R. Hursh, 2014). The essential value (*EV*), conceptualized as a strength of the reinforcer to maintain operant behavior in the face of rising behavioral cost, was derived from the economic demand model. Additional parameters derived from the economic demand model were used in supplemental analyses (*Q_0_*, *α*, *P_max_*, and *O_max_*). All linear mixed-effects analyses were performed in R 3.6.1 (R Core Team, 2019) using {nlme} package (Pinheiro et al., 2017). The linear mixed-effects modeling approach provides a number of advantages when compared to ANOVAs. For example, this analysis does not require the assumption that the relation between the covariate and the outcome is the same across the groups and thus does not require meeting the assumption of homogeneity. Furthermore, unlike ANOVA, linear mixed-effects modeling does not assume that the different cases of data were independent and hence can model relations between different outcomes, which may be interrelated. Linear mixed-effects modeling is also more robust in dealing with missing data or unequal group sizes which is often the case in preclinical animal models. For these reasons, most of the effects in this study were analyzed using linear mixed-effects modeling. Pairwise comparisons were performed using estimates from the model. Linear regression analysis and least-squares nonlinear fit for the assessment of economic demand parameters were performed using GraphPad Prism version 8.2.1 (GraphPad Software, Inc., La Jolla, CA).

##### 2.7.1.3. Experiment 2

The amount of sucrose consumed was used as a primary dependent measure. Sucrose consumption was assessed using daily consumption as a repeated measure and using total consumption as a supplementary measure.

## 3. Results

### 3.1. Experiment 1

#### 3.1.1. Economic demand for sucrose

Table 1 summarizes all the parameters derived from the behavioral economics model by varenicline condition and nicotine dose. Table 2 summarizes parameters from statistical tests involving *EV* parameter, our primary dependent measure, that was derived from the economic demand equation. Omnibus assessment of the effects of varenicline and nicotine treatments on the *EV* revealed a significant effect of Nicotine Dose, no effect of Varenicline Condition, and no interaction (see Table 2). To better understand the effect of Nicotine Dose, the *EV* was then analyzed separately within each Varenicline Condition. There was a significant effect of Nicotine Dose on *EV* of rats treated with 0.0 and 0.1 mg/kg of varenicline (see Table 2). Pairwise comparisons of Nicotine Dose (compared to 0.0 mg/kg) within each Varenicline Condition showed that 0.4 mg/kg nicotine treatment significantly decreased economic demand in rats not treated with varenicline (0.0 mg/kg Varenicline Condition; *b* = −10.40*, t*(22) = −4.40*, p* = 0.0002) and in rats treated with low varenicline dose (0.1 mg/kg Varenicline Condition; *b* = −4.38*, t*(22) = −3.12*, p* = 0.0049; Figure 2A).

**Table 1.** Major parameters derived from the behavioral economics model.

| | | $EV$ | | | $Q_o$ | | | $\alpha$ | | | $O_{max}$ | | | $P_{max}$ | | |
| --- | --- | --- | --- | --- | --- | --- | --- | --- | --- | --- | --- | --- | --- | --- | --- | --- |
| Nicotine Dose (mg/kg) → |  | 0.0 | 0.1 | 0.4 | 0.0 | 0.1 | 0.4 | 0.0 | 0.1 | 0.4 | 0.0 | 0.1 | 0.4 | 0.0 | 0.1 | 0.4 |
| Condition | $\kappa$ | | | | | | | | | | | | | | | |
| Var 0.0 | 2.5 | 54.14 | 52.02 | **43.74** | 197.95 | 211.00 | **218.35** | 5.11E-05 | 5.24E-05 | **6.15E-05** | 1510.83 | 1451.67 | **1220.46** | 24.13 | 21.70 | **17.46** |
| Var 0.1 | 2.5 | 50.67 | 47.79 | **46.28** | 203.46 | 217.98 | **223.05** | 6.26E-05 | 6.78E-05 | 6.79E-05 | 1413.99 | 1333.63 | **1291.48** | 21.61 | 19.44 | **18.90** |
| Var 1.0 | 2.5 | 59.30 | 58.25 | 54.40 | 211.82 | 215.57 | 217.94 | 5.28E-05 | 5.16E-05 | 5.72E-05 | 1654.82 | 1625.58 | 1518.13 | 25.23 | 24.38 | 22.38 |

Numbers in **bold** indicate significantly different responding ( $p < 0.05$ ) from responding observed at 0.0 mg/kg nicotine dose.

**Table 2.** The statistical output from analyses assessing mixed-effects of varenicline group, nicotine dose, and their interaction on *essential value* derived from the economics demand model.

|  | Model | df | AIC | BIC | logLik | Test | L.Ratio | p-value |
| --- | --- | --- | --- | --- | --- | --- | --- | --- |
| **Omnibus Assessment** |  |  |  |  |  |  |  |  |
| *Baseline Model (DV=EV)* | 1 | 4 | 844.511 | 855.2397 | -418.256 |  |  |  |
| Adding Var Factor | 2 | 6 | 846.782 | 862.8747 | -417.391 | 1 vs 2 | 1.72924 | 0.4212 |
| Adding Nic Factor | 3 | 8 | 832.331 | 853.788 | -408.166 | 2 vs 3 | 18.451 | **0.0001** |
| Adding Var × Nic | 4 | 12 | 835.233 | 867.4185 | -405.616 | 3 vs 4 | 5.09808 | 0.2774 |
| **Varenicline 0.0 condition** |  |  |  |  |  |  |  |  |
| *Baseline Model (DV=EV)* | 1 | 4 | 283.115 | 289.4491 | -137.558 |  |  |  |
| Adding Nic Factor | 2 | 6 | 270.653 | 280.154 | -129.326 | 1 vs 2 | 16.4622 | **0.0003** |
| **Varenicline 0.1 condition** |  |  |  |  |  |  |  |  |
| *Baseline Model (DV=EV)* | 1 | 4 | 259.082 | 265.4163 | -125.541 |  |  |  |
| Adding Nic Factor | 2 | 6 | 254.015 | 263.5158 | -121.007 | 1 vs 2 | 9.06753 | **0.0107** |
| **Varenicline 1.0 condition** |  |  |  |  |  |  |  |  |
| *Baseline Model (DV=EV)* | 1 | 4 | 299.833 | 306.1671 | -145.917 |  |  |  |
| Adding Nic Factor | 2 | 6 | 301.551 | 311.052 | -144.776 | 1 vs 2 | 2.28217 | 0.3195 |

Numbers in **bold** indicate statistical significance ( $p < 0.05$ ).

**Figure 2:**
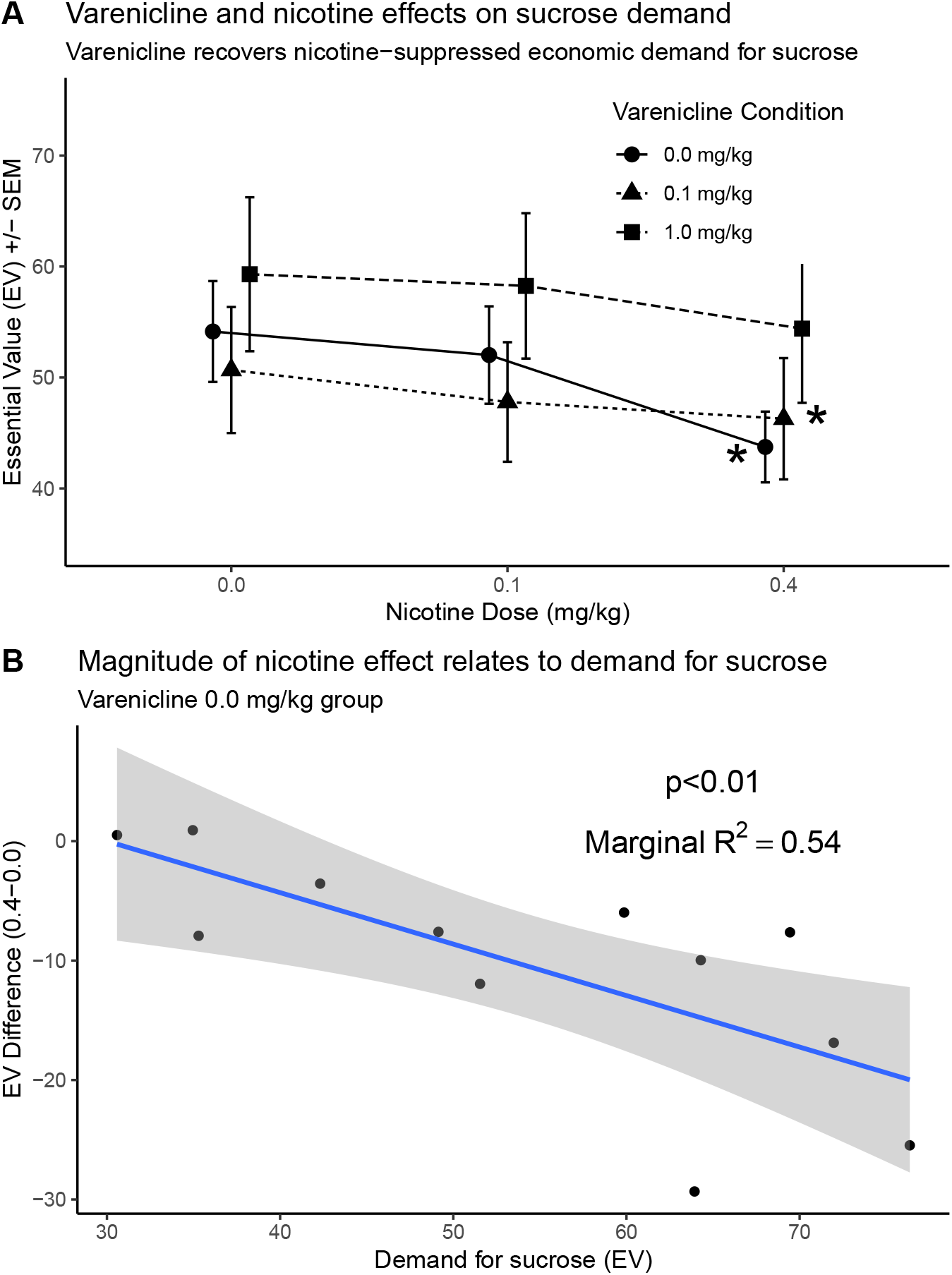
(A) *EV* for each Varenicline Condition across nicotine doses that was derived from the economic demand model of sucrose consumption. * Indicates significantly lower *EV* in comparison to a corresponding 0.0 mg/kg nicotine dose. (B) Individual demand for sucrose explains 54 % of the variance in the magnitude of nicotine-induced decrease in *EV* that is visualized as *EV* difference score.

Supplemental analyses of additional economic demand parameters (*Q_0_*, *α*, *P_max_*, and *O_max_*) revealed a significant effect of Nicotine Dose, no effect of Varenicline Condition, and no interaction across all the parameters (Table S1; data not shown). The analyses of those parameters for each Varenicline Condition and the associated pairwise nicotine dose comparisons are presented in Figure S2 (data not shown). The assessment of these additional parameters showed that nicotine increased elasticity of nicotine demand (*α*), decreased *P_max_*, decreased *O_max_*, and surprisingly increased responding when the price of a reinforcer is conceptualized as zero or free (*Q_0_*; Tables 1 and S2). In the assessment of these effects, it is important to note that *α* has an inverse relationship to *EV* and that *P_max_* (the price at the point of maximal responding) closely relates to *O_max_* (maximum responses observed at *P_max_*) and that both of them often correlate to *EV*. Therefore, the changes in *α*, *P_max_*, and *O_max_* largely parallel changes in *EV*. For these reasons, further discussion will center around the effects associated with *EV* and *Q_0_*.

We then assessed how the magnitude of the nicotine effect on the demand for sucrose related to the individual economic demand for sucrose. We showed above that nicotine (0.4 mg/kg) significantly decreased the *essential value* of sucrose in conditions where rats received no or low dose of varenicline and there was evidence of individual variability underlying this effect. We hypothesized that rats with higher economic demand for sucrose would show a more pronounced response to the nicotine treatment. To assess this assumption, we subtracted the *EV* observed after treatment with 0.4 mg/kg of nicotine from the *EV* after pretreatment with 0.0 mg/kg of nicotine (*EV* (0.4)-*EV* (0.0); negative numbers indicate decrease in *EV*) and compared this difference to the *EV* of sucrose in the absence of nicotine treatment ([*EV* (0.4)-*EV* (0.0)] vs *EV* (0.0); *κ* constrained to 2.5). We performed this analysis for each Varenicline Condition. We detected the significant relationship between the magnitude of nicotine effect (*EV* difference) and the individual demand for sucrose (*EV*) only in rats that have not been treated with varenicline (0.0 mg/kg Varenicline Condition (*χ*^2^(1) = 8.93*, p* = 0.002). These results show that the *EV* of sucrose can explain 54 % of the variance (marginal *R*^2^ = 0.54) in the magnitude of response (decrease in *EV*) to nicotine treatment (Figure 2B). These results show that nicotine decreased the *EV* of sucrose. The magnitude of nicotine effect on the demand for sucrose (*EV*) in rats not treated with varenicline (0.0 mg/kg condition) was dependent on the individual demand for sucrose, where rats with higher demand for sucrose showed a higher magnitude of decrease. On the other hand, varenicline pretreatment blocked the effect of nicotine on the economic demand for sucrose.

#### 3.1.2. Responding on PR schedule of reinforcement tests

Responding on the PR schedule of reinforcement was assessed separately for each Varenicline Condition (Figure 3). The statistical output from these analyses is presented in Table 3. For each Varenicline Dose we built a model with Varenicline Condition, Nicotine Dose, and their interaction as predictors. We analyzed data from PR schedule of reinforcement tests by Varenicline dose because it allows to compare the effect of nicotine on responding for sucrose across each history with varenicline treatment (Varenicline Condition). With this in mind, the omnibus assessment of main effects for PR tests involving 0.0 mg/kg varenicline treatment revealed a significant effect of Nicotine Dose, no effect of Varenicline Condition, although it was trending towards significance with *p* = 0.0606, and no interaction. The trending effect of Varenicline Condition was followed by pairwise comparisons. Overall, the responding of rats with a history of 1.0 mg/kg varenicline treatment was significantly lower than responding of rats that were not treated with varenicline prior to PR tests (0.0 mg/kg; *b* = −2.06*, t*(33) = −2.17*, p* = 0.0372; see # symbol in Figure 3A). The responding of rats with a history of 0.1 mg/kg varenicline treatment was trending towards being significantly lower than the responding of rats that were not treated with varenicline prior to PR tests (0.0 mg/kg; *b* = −1.94*, t*(33) = −2.03*, p* = 0.0500; Figure 3A). Follow-up assessments of the effect of nicotine for each Varenicline Condition showed a significant effect of Nicotine Dose within each condition and significantly lower responding for sucrose after pretreatment with 0.4 mg/kg of nicotine (Table 3; see * symbols in Figure 3A).

**Table 3.** Statistical output from the progressive ratio schedule of reinforcement tests by history of varenicline condition (Varenicline Condition). For the nicotine dose pairwise comparisons, the model was built with Nicotine Dose as a single predictor and the estimates for comparisons were derived from that model.

|  | Figure | Model | df | AIC | BIC | logLik | Test | L.Ratio | p-value | Nicotine dose (mg/kg) comparison |  |
| --- | --- | --- | --- | --- | --- | --- | --- | --- | --- | --- | --- |
|  |  |  |  |  |  |  |  |  |  | 0.1 vs 0.0 | 0.4 vs 0.0 |
| **Var 0.0 treatment \| Condition×Nic** | **3A** |  |  |  |  |  |  |  |  |  |  |
| *Baseline Model (DV=Sucrose Earned)* |  | 1 | 4 | 534.07 | 544.80 | -263.04 |  |  |  |  |  |
| Adding Varenicline Condition Factor |  | 2 | 6 | 532.47 | 548.56 | -260.23 | 1 vs 2 | 5.61 | 0.0606 |  |  |
| Adding Nicotine Dose Factor |  | 3 | 8 | 499.74 | 521.20 | -241.87 | 2 vs 3 | 36.73 | <.0001 |  |  |
| Adding Condition × Nic |  | 4 | 12 | 506.02 | 538.20 | -241.01 | 3 vs 4 | 1.73 | 0.7861 |  |  |
| **Var 0.0 treatment \| Var 0.0 Condition** | **3A●** |  |  |  |  |  |  |  |  |  |  |
| *Baseline Model (DV=Sucrose Earned)* |  | 1 | 4 | 170.69 | 177.13 | -81.34 |  |  |  |  |  |
| Adding Nicotine Dose Factor |  | 2 | 6 | 160.78 | 170.44 | -74.39 | 1 vs 2 | 13.91 | 0.0010 | b = -0.11, t(22) = -0.19, p = 0.8496 | b = -2.20, t(22) = -3.72, p = 0.0012 |
| **Var 0.0 treatment \| Var 0.1 Condition** | **3A▲** |  |  |  |  |  |  |  |  |  |  |
| *Baseline Model (DV=Sucrose Earned)* |  | 1 | 4 | 188.50 | 194.72 | -90.25 |  |  |  |  |  |
| Adding Nicotine Dose Factor |  | 2 | 6 | 181.08 | 190.41 | -84.54 | 1 vs 2 | 11.42 | 0.0033 | b = 0.75, t(21) = -0.83, p = 0.4129 | b = -2.48, t(21) = -2.68, p = 0.0141 |
| **Var 0.0 treatment \| Var 1.0 Condition** | **3A■** |  |  |  |  |  |  |  |  |  |  |
| *Baseline Model (DV=Sucrose Earned)* |  | 1 | 4 | 180.02 | 186.35 | -86.01 |  |  |  |  |  |
| Adding Nicotine Dose Factor |  | 2 | 6 | 169.88 | 179.38 | -78.94 | 1 vs 2 | 14.14 | 0.0009 | b = -0.17, t(22) = -0.25, p = 0.8045 | b = -2.50, t(22) = -3.75, p = 0.0011 |
| **Var 0.1 treatment \| Condition×Nic** | **3B** |  |  |  |  |  |  |  |  |  |  |
| *Baseline Model (DV=Sucrose Earned)* |  | 1 | 4 | 484.72 | 495.33 | -238.36 |  |  |  |  |  |
| Adding Varenicline Condition Factor |  | 2 | 6 | 485.15 | 501.07 | -236.57 | 1 vs 2 | 3.57 | 0.1680 |  |  |
| Adding Nicotine Dose Factor |  | 3 | 8 | 457.34 | 478.57 | -220.67 | 2 vs 3 | 31.81 | <.0001 |  |  |
| Adding Condition × Nic |  | 4 | 12 | 462.72 | 494.57 | -219.36 | 3 vs 4 | 2.62 | 0.6226 |  |  |
| **Var 0.1 treatment \| Var 0.0 Condition** | **3B●** |  |  |  |  |  |  |  |  |  |  |
| *Baseline Model (DV=Sucrose Earned)* |  | 1 | 4 | 160.37 | 166.47 | -76.18 |  |  |  |  |  |
| Adding Nicotine Dose Factor |  | 2 | 6 | 148.67 | 157.83 | -68.33 | 1 vs 2 | 15.70 | 0.0004 | b = -0.13, t(20) = -0.23, p = 0.8175 | b = -2.17, t(20) = -4.14, p = 0.0005 |
| **Var 0.1 treatment \| Var 0.1 Condition** | **3B▲** |  |  |  |  |  |  |  |  |  |  |
| *Baseline Model (DV=Sucrose Earned)* |  | 1 | 4 | 164.32 | 170.66 | -78.16 |  |  |  |  |  |
| Adding Nicotine Dose Factor |  | 2 | 6 | 161.82 | 171.32 | -74.91 | 1 vs 2 | 6.51 | 0.0386 | b = -0.17, t(22) = -0.30, p = 0.7668 | b = -1.33, t(22) = -2.40, p = 0.0252 |
| **Var 0.1 treatment \| Var 1.0 Condition** | **3B■** |  |  |  |  |  |  |  |  |  |  |
| *Baseline Model (DV=Sucrose Earned)* |  | 1 | 4 | 171.01 | 177.23 | -81.51 |  |  |  |  |  |
| Adding Nicotine Dose Factor |  | 2 | 6 | 163.34 | 172.67 | -75.67 | 1 vs 2 | 11.67 | 0.0029 | b = -0.83, t(21) = -1.32, p = 0.2002 | b = -2.25, t(21) = -3.69, p = 0.0014 |
| **Var 1.0 treatment \| Condition×Nic** | **3C** |  |  |  |  |  |  |  |  |  |  |
| *Baseline Model (DV=Sucrose Earned)* |  | 1 | 4 | 501.15 | 511.95 | -246.57 |  |  |  |  |  |
| Adding Varenicline Condition Factor |  | 2 | 6 | 498.64 | 514.84 | -243.32 | 1 vs 2 | 6.51 | 0.0385 |  |  |
| Adding Nicotine Dose Factor |  | 3 | 8 | 477.64 | 499.24 | -230.82 | 2 vs 3 | 25.00 | <.0001 |  |  |
| Adding Condition × Nic |  | 4 | 12 | 483.94 | 516.34 | -229.97 | 3 vs 4 | 1.70 | 0.7906 |  |  |
| **Var 1.0 treatment \| Var 0.0 Condition** | **3C●** |  |  |  |  |  |  |  |  |  |  |
| *Baseline Model (DV=Sucrose Earned)* |  | 1 | 4 | 152.70 | 159.03 | -72.35 |  |  |  |  |  |
| Adding Nicotine Dose Factor |  | 2 | 6 | 146.26 | 155.76 | -67.13 | 1 vs 2 | 10.44 | 0.0054 | b = -0.35, t(19) = -0.73, p = 0.4720 | b = -1.62, t(19) = -3.24, p = 0.0043 |
| **Var 1.0 treatment \| Var 0.1 Condition** | **3C▲** |  |  |  |  |  |  |  |  |  |  |
| *Baseline Model (DV=Sucrose Earned)* |  | 1 | 3 | 175.84 | 180.67 | -84.92 |  |  |  |  |  |
| Adding Nicotine Dose Factor |  | 2 | 5 | 168.49 | 176.55 | -79.25 | 1 vs 2 | 11.35 | 0.0034 | b = -0.91, t(23) = -1.32, p = 0.1974 | b = -2.45, t(23) = -3.61, p = 0.0015 |
| **Var 1.0 treatment \| Var 1.0 Condition** | **3C■** |  |  |  |  |  |  |  |  |  |  |
| *Baseline Model (DV=Sucrose Earned)* |  | 1 | 4 | 176.35 | 182.79 | -84.17 |  |  |  |  |  |
| Adding Nicotine Dose Factor |  | 2 | 6 | 174.69 | 184.36 | -81.35 | 1 vs 2 | 5.66 | 0.0591 | b = -0.20, t(22) = -0.30, p = 0.7663 | b = -1.50, t(22) = -2.23, p = 0.0362 |

Numbers in **bold** indicate statistical significance (p<0.05).

**Figure 3:**
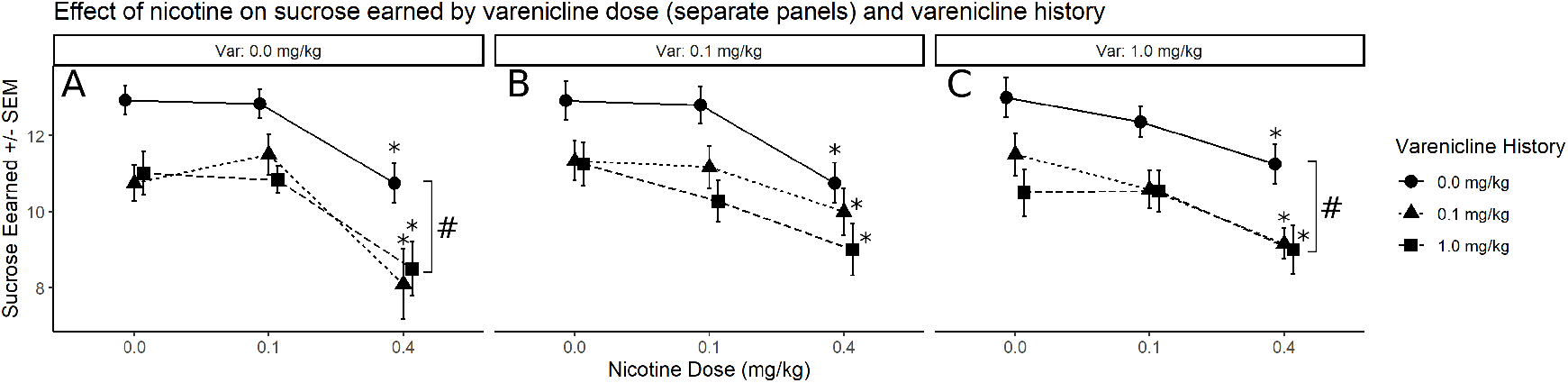
Sucrose earned after pretreatment with a combination of varenicline (0.0, 0.1, or 1.0 mg/kg; separate panels A-C) and nicotine (0.0, 0.1, 0.4 mg/kg). The analyses and visualization by Varenicline Dose allow to compare the effect of nicotine on responding for sucrose across each history of varenicline treatment (Varenicline History plotted as separate lines). * Indicates the significantly lower number of sucrose presentations earned in comparison to a corresponding 0.0 mg/kg nicotine dose. # Indicates significant difference in sucrose earned across all nicotine doses.

The omnibus assessment of main effects for PR tests involving 0.1 mg/kg varenicline treatment revealed a significant effect of Nicotine Dose, no effect of Varenicline Condition, and no interaction. Follow-up assessments of the effect of nicotine for each Varenicline Condition showed a significant effect of Nicotine Dose within each condition and significantly lower responding for sucrose after pretreatment with 0.4 mg/kg of nicotine (Table 3; see * symbols in Figure 3B).

The omnibus assessment of main effects for PR tests involving 1.0 mg/kg varenicline treatment revealed significant effects of Nicotine Dose, Varenicline Condition, and no interaction. Overall, the responding of rats with a history of 1.0 mg/kg varenicline treatment was significantly lower than the responding of rats that were not treated with varenicline prior to PR tests (0.0 mg/kg; *b* = 0.92*, t*(33) = −2.48*, p* = 0.0184; see # symbol in Figure 3C). The responding of rats with a history of 0.1 mg/kg varenicline treatment was trending towards being significantly lower than the responding of rats that were not treated with varenicline prior to PR tests (0.0 mg/kg; *b* = 0.92*, t*(33) = −2.02*, p* = 0.0515; Figure 3C). Follow-up assessments of the effect of nicotine for each Varenicline Condition showed a significant effect of Nicotine Dose within each condition and significantly lower responding for sucrose after pretreatment with 0.4 mg/kg of nicotine (Table 3; see * symbols in Figure 3A).

These results show that the history of varenicline treatment is a predictor of responding for sucrose on a PR schedule of reinforcement. Specifically, rats with a history of varenicline treatment responded significantly lower for sucrose across nicotine doses than rats that have not being exposed to varenicline prior to the PR tests. Finally, consistent with the results from the economic demand phase of the experiment, nicotine (0.4 mg/kg) decreased responding for sucrose independent on varenicline treatment or previous history with varenicline.

### 3.2. Experiment 2

The assessment of sucrose consumption over 25 daily sessions revealed significant effects of Condition (*χ*^2^(3) = 11.51*, p* = 0.0093), Day (*χ*^2^(27) = 1119.80*, p* < 0.0001), and their interaction (*χ*^2^(99) = 159.27*, p* < 0.0001). Overall, the consumption of sucrose after pretreatment with nicotine (0.4 mg/kg; Sal-Nic condition) was significantly lower than no treatment controls (Sal-Sal condition; Figure 4A). A complimentary assessment of total consumption over the course of 25 daily sessions revealed a significant effect of Condition (*χ*^2^(3) = 11.37*, p* = 0.0099) and pairwise comparisons showed that the total consumption of sucrose after pretreatment with nicotine (0.4 mg/kg; Sal-Nic condition) was significantly lower than the total consumption of no-treatment controls (Sal-Sal condition; Figure 4B).

**Figure 4:**
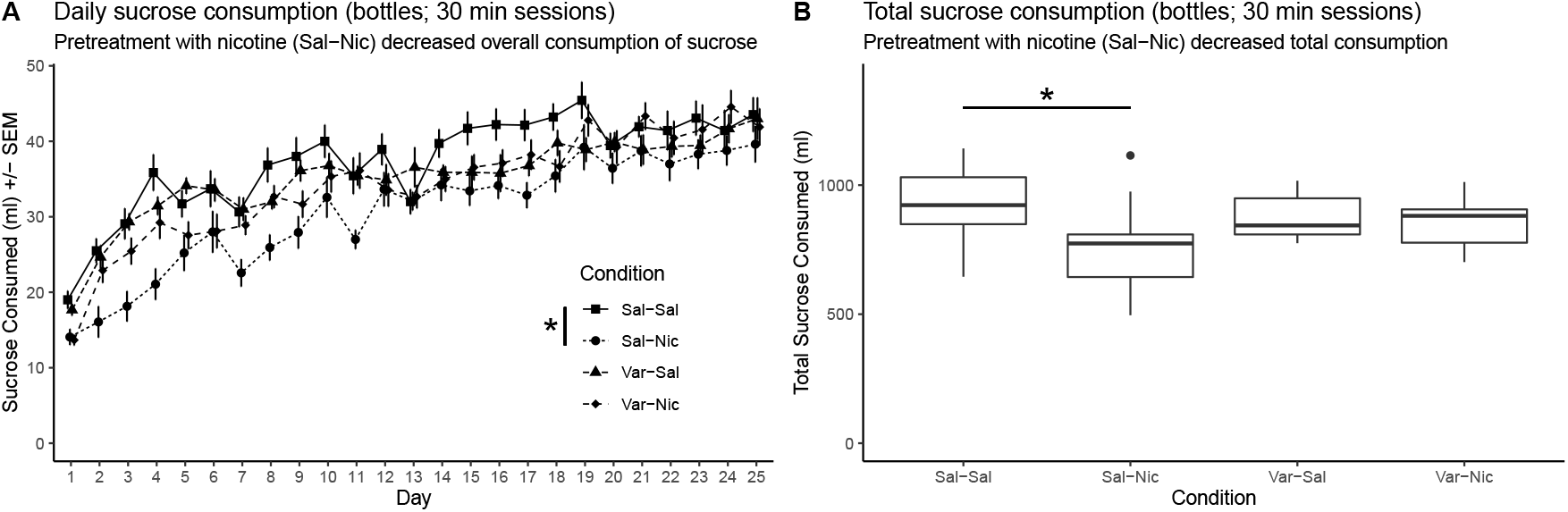
Rats were assigned to conditions using a 2 2 design with varenicline (0.0 or 1.0 mg/kg; identified on the figure as Sal or Var respectively) and nicotine (0.0 or 0.4 mg/kg; identified on the figure as Sal or Nic respectively) as between-subjects variables. (A) Daily sucrose consumption. (B) Total sucrose consumption.

## 4. Discussion

Varenicline is a widely prescribed medication used to treat nicotine dependence. Varenicline is often prescribed in conjunction with nicotine use in the early phase of the treatment regiment (Pfizer, 2020; Tonstad et al., 2006). Although it is unclear how effective varenicline is in improving long-term abstinence from nicotine use (Hajek et al., 2009; Lancaster et al., 2006), it remains one of the most prescribed medications for smoking cessation around the world, and it has been shown to be more efficacious than bupropion in improving short-term abstinence (Cahill et al., 2013; Gonzales, 2006; Jorenby et al., 2006). The precise mechanism underlying varenicline effects on nicotine use is not fully understood, but the existing evidence suggests that its ability to mimic and substitute for the effects of nicotine—by virtue of a similar pharmacological profile—likely involved in its ability to reduce cravings and attenuate the reinforcing effects of nicotine. One of the important considerations when prescribing substance cessation treatment is the effect of that treatment on other reinforcers in the environment and food reinforcement in particular. Decreasing the reinforcing value of other reinforcers in the environment may result in unanticipated negative effects like reduced compliance with the treatment protocol or reduced rates of long-term abstinence. With that in mind, the effects of varenicline alone or in conjunction with nicotine on the reinforcing effects of foods are underexplored. The present study was designed to address this gap by testing the effects of varenicline and nicotine on the economic demand for sucrose. We found that a) nicotine alone decreased the *essential value* of sucrose, b) the magnitude of nicotine-induced reduction in demand for sucrose in rats not treated with varenicline was dependent on baseline individual demand for sucrose c) varenicline rescued nicotine-induced reduction in economic demand for sucrose, and d) history of varenicline treatment predicted responding for sucrose on a PR schedule of reinforcement. Specifically, rats with a history of varenicline treatment responded significantly lower for sucrose across nicotine doses than rats that have not being exposed to varenicline prior to the PR tests. In addition, we showed that nicotine also decreased the consumption of sucrose in home cages and that varenicline also rescued this effect. Overall, these findings suggest that the effects of varenicline and nicotine interact to affect motivation for appetitive rewards. Importantly, we show for the first time that varenicline has a long-term effect on food reinforcement—the effect that is essentially unexplored for its role in smoking cessation.

Previous studies firmly established that nicotine increases rewarding effects of other reinforcers in the environment, environmental cues, and increases incentive value of other non-drug reinforcers (Caggiula et al., 2009; Grimm et al., 2012; Palmatier et al., 2013; Perkins and Karelitz, 2013). However, how nicotine affects the reinforcing effects of food reinforcement is not fully understood because of existing conflicting reports in the literature. For example, some reports showed that contingent or non-contingent nicotine administration increases sucrose intake (Grimm et al., 2012; Jias and Ellison, 1990; Smith and Roberts, 1995), whereas other reports show that nicotine self-administration or exposure to cigarette smoke decrease sucrose and food intake (Bunney et al., 2016; Chen et al., 2005). These reports cited above used different approaches to assess the effects of nicotine on food or sucrose consumption ranging from measuring daily chow consumption, sucrose consumption in home cages, or self-administration of chow or sucrose pellets no name a few. On the other hand, the effects of nicotine on food or sucrose reinforcement have not been examined using a behavioral economics approach which provides a more comprehensive assessment of consumption expenditure over a range of workload demands. Using this reinforcer demand modeling, our findings support previous reports showing that nicotine decreases sucrose intake. We also extend previous findings by demonstrating that nicotine decreases the *essential value* of sucrose and that the magnitude of this effect relates to baseline individual demand for sucrose. Specifically, we show that individual demand for sucrose predicts response to nicotine, where rats with higher demand for nicotine are more sensitive to the effects of nicotine and show the higher magnitude of decrease in demand for nicotine. On the other hand, nicotine increased responding for sucrose at modeled “zero” price, which was evident by the increase in *Q_0_* value (see Table 1, varenicline 0.0 mg/kg condition). This differential effect on reinforcing effects of sucrose suggests that the “cost” of access to reinforcement may play a role in nicotine-modulated effects on motivation for reward. To better understand the effects observed in Experiment 1, we conducted Experiment 2 that was designed to assess the effect of nicotine on sucrose consumption in home cages. Thus, Experiment 2 provides an alternative methodology to assess and confirm findings from Experiment 1. Our findings from Experiment 2 show that nicotine also reduced sucrose consumption in home cages. This nicotine-induced reduction in home cage sucrose consumption parallels the effect of nicotine on *EV* observed in Experiment 1. Conversely, this nicotine-induced reduction in home cage sucrose consumption is contrary to what one may expect after the observed effects of nicotine on the *Q_0_* of sucrose consumption in Experiment 1. In Experiment 1 we show that nicotine increased *Q_0_* of responding for sucrose. *Q_0_* is conceptualized as a responding observed at the price of zero (simulated free cost), and one may expect that this should translate to the effect of nicotine on the consumption of sucrose in home cages where the “price” of access to sucrose can be conceptualized as being relatively low or close to “free” (some energy expenditure still required to access bottle spout). Overall, these effects demonstrate that nicotine decreased economic demand for sucrose, conceptualized here as a decrease in *essential value*, and decreased consumption of sucrose in home cages. Surprisingly, nicotine also increased the reinforcing value of sucrose at modeled “free” cost, indicating a possible role of cost to access a reinforcer in the effects of nicotine on reward and reinforcement.

Varenicline is one of the most effective pharmacotherapies approved for smoking cessation. However, long-term abstinence rates following the treatment with varenicline are relatively low and vary across studies (Cahill et al., 2013; Jordan and Xi, 2018; Rosen et al., 2018). One of the distinguishing features of varenicline treatment is its concomitant administration with nicotine use. While there are studies investigating the effects of nicotine or varenicline alone on motivation for primary reinforcers, the interaction effect of these substances on primary reinforcement is not fully understood. Existing research shows that primary reinforcing effects of varenicline are weak but varenicline can be self-administered in preclinical settings under low workload requirements (Paterson et al., 2010; Rollema et al., 2007; Schassburger et al., 2015). Varenicline effects on motivation for other reinforcers are complex and range from attenuation of nicotine self-administration using short-access protocols (George et al., 2011; O’Connor et al., 2010; Rollema et al., 2007), no effect on nicotine self-administration using long-access protocols (Kazan et al., 2020), enhancement of motivation for sensory reinforcement (Barrett et al., 2018; Levin et al., 2012), and antagonism of reinforcement-enhancing effects of nicotine (Levin et al., 2012) to name a few. Our study here further expands our understanding of varenicline effects on motivation for primary reinforcers. In this study, we show that varenicline has no effect on the economic demand for liquid sucrose. Because sucrose is a potent primary reinforcer, it is often difficult to assess the effects of other stimuli on motivation for sucrose. Rats will often demonstrate a ceiling effect (maximum allowed responding) when responding for sucrose is assessed using low or moderate workload requirements. Our approach, on the other hand, samples behavior across a range of workload requirements and then further interprets this responding using a well-established behavioral economics model. To further expand our understanding of varenicline effects on motivation for primary reinforcers a more parametric approach including a range of primary reinforcers and length of access may be required.

The effectiveness of varenicline as a smoking cessation treatment is often attributed to its agonistic effects at nicotinic acetylcholine receptors and its ability to induce dopamine in the mesolimbic system (Coe et al., 2005; Ericson et al., 2009; Rollema et al., 2007). The mechanism by which nicotine enhances dopamine levels in the mesolimbic system involve agonistic actions at nicotinic acetylcholine receptors in the ventral tegmental area (De Biasi and Dani, 2011). Interestingly, previous studies show that pretreatment with varenicline reduces nicotine-evoked dopamine release in nucleus accumbens and suggest that varenicline interaction at nicotinic acetylcholine receptors in the ventral tegmental area may by central to this effect (Ericson et al., 2009; Rollema et al., 2007). Our study shows that varenicline rescued nicotine-induced decrease in the *essential value* of sucrose. Although our study here does not provide an insight into neural mechanisms underlying this effect, we can only speculate based on previous reports that varenicline interactions with nicotine in the mesolimbic system may be involved in the blockade of nicotine-induced decrease in the *essential value* of sucrose. Furthermore, it is possible that the combined summative effects of the reinforcing effects of nicotine and of sucrose are behaviorally limited by a reinforcement ceiling, which varenicline attenuates by antagonizing the reinforcing effects of nicotine.

One of the most interesting effects observed in our study is the long-term effect of varenicline treatment on responding for sucrose during PR schedule of reinforcement tests. We show that rats with a history of varenicline treatment responded significantly less for sucrose on PR schedule of reinforcement than rats that have not been exposed to varenicline in the previous phase of the study. This diminished reduction in responding for sucrose suggests a long-term effect of varenicline treatment on the reward system. Previous studies indicate that nicotinic acetylcholine receptors desensitize after stimulation with nicotine and up-regulate following long-term exposure to nicotine (Benwell et al., 1995; Collins et al., 1994; Schwartz and Kellar, 1985). Likewise, the varenicline desensitizes nicotinic acetylcholine receptors (Rollema et al., 2010) and this effect may explain long-term effects we observed in our study. Furthermore, repeated treatment with varenicline decreases dopamine levels in nucleus accumbens evoked by concomitant administration of nicotine and ethanol, while acute treatment with varenicline did not, suggesting a possible role of desensitization of nicotinic receptors in this effect (Ericson et al., 2009). In our study, all rats have been exposed to nicotine during the demand phase of the experiment preceding testing on PR schedule of reinforcement. During that demand phase of the study, some rats were exposed to varenicline (0.1 and 1.0 mg/kg) on a daily basis, while control group (0.0 mg/kg) was not. During the PR tests responding of rats in both conditions with previous history of varenicline treatment was equally lower than responding of rats that have not been previously treated with varenicline. Our findings suggest that the effects of varenicline on the reward system may be distinct from the effects of nicotine. Although one may argue that it is possible that the reduction of responding for sucrose during the PR schedule of reinforcement tests represents the additive effect of nicotine and varenicline on the reward system, the fact that rats in both low and high dose varenicline conditions responded equally during those tests suggests otherwise. Additional studies will be required to further understand the nature of these effects.

## 5. Acknowledgments

S. Charntikov was supported by GM113131 (CIBBR, P20) while preparing this manuscript for publication.

---

**Table S1.**
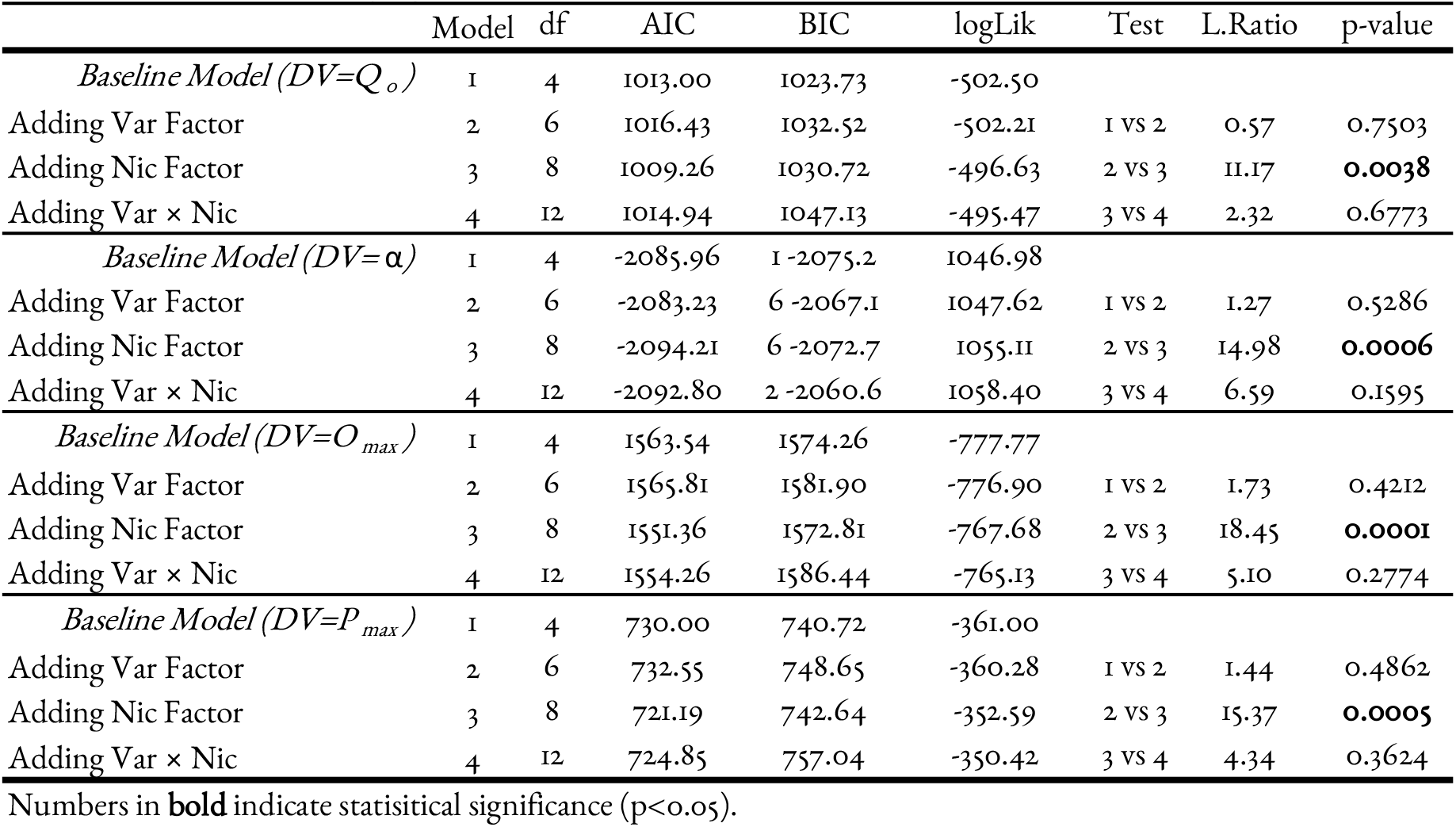
The statistical output from analyses assessing mixed effects of varenicline group, nicotine dose, and their interaction for additional major parameters derived from the economics demand model.

**Figure S2.**
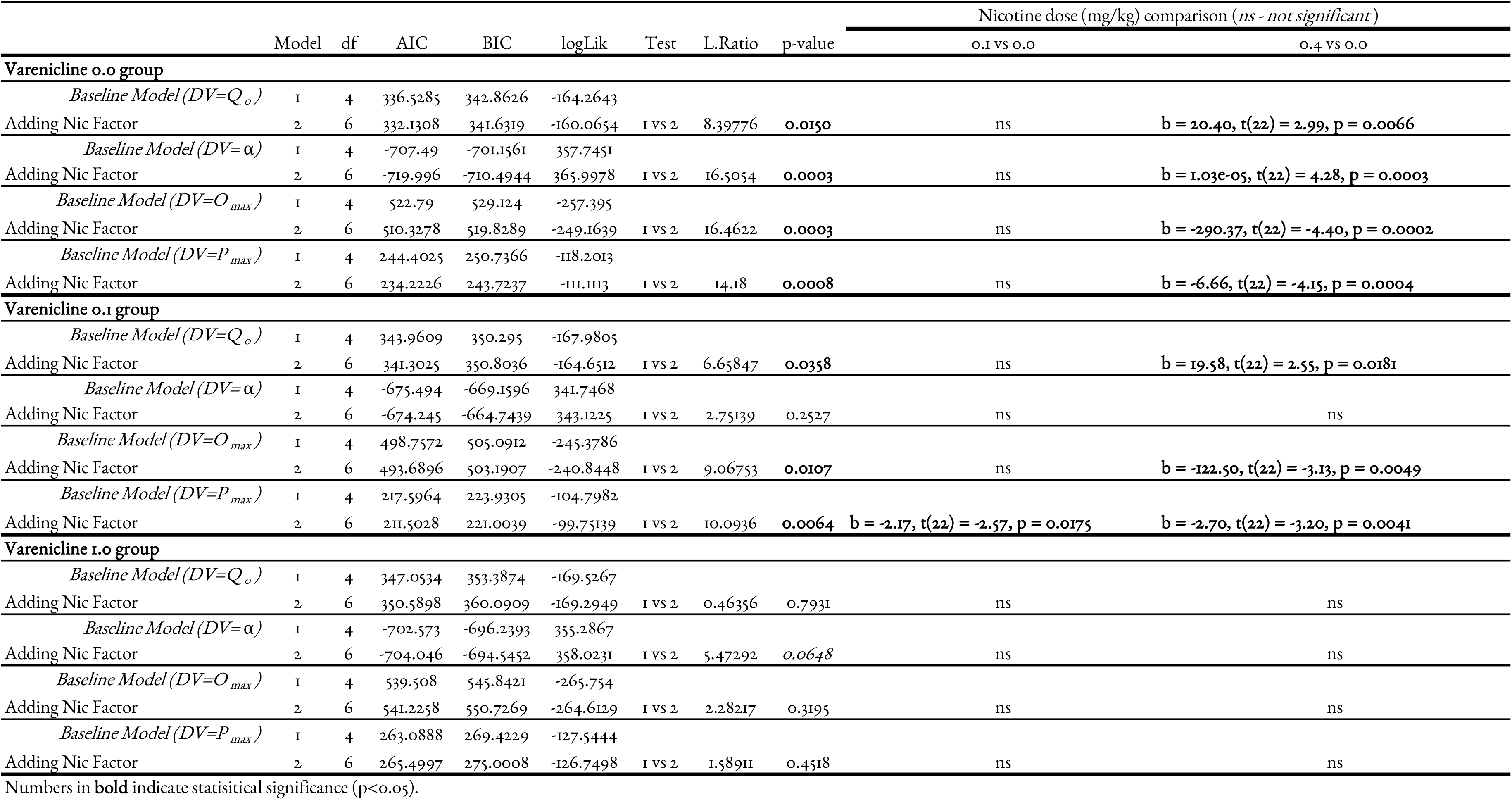
The statistical output from analyses assessing effects of nicotine on additional parameters derived from behavioral economics model for each varenicline condition and associated pairwise comparisons.

